# SMART: reference-free deconvolution for spatial transcriptomics using marker-gene-assisted topic models

**DOI:** 10.1101/2023.06.20.545793

**Authors:** C Yang, DD Sin, RT Ng

## Abstract

Spatial transcriptomics (ST) offers valuable insights into gene expression patterns within the spatial context of tissue. However, most technologies do not have a single-cell resolution, masking the signal of the individual cell types. Here, we present SMART, a reference-free deconvolution method that simultaneously infers the cell type-specific gene expression profile and the cellular composition at each spot. Unlike most existing methods that rely on having a single-cell RNA-sequencing dataset as the reference, SMART only uses marker gene symbols as the prior knowledge to guide the deconvolution process and outperforms the existing methods in realistic settings when an ideal reference dataset is unavailable. SMART also provides a two-stage approach to enhance its performance on cell subtypes. Allowing the inclusion of covariates, SMART provides condition-specific estimates and enables the identification of cell type-specific differentially expressed genes across conditions, which elucidates biological changes at a single-cell-type resolution.

## Introduction

Spatial transcriptomics (ST) is a cutting-edge technology that enables scientists to measure gene expression patterns across different tissue regions with spatial information^1–3^. In general, with the current ST platforms, the measured spots on a tissue sample do not demonstrate single-cell resolutions but contain a complex mixture of multiple cell types^4–6^. This hinders our understanding of the spatial organization of these cells and precludes the identification of cell type-specific transcriptomic signatures^7^. Understanding the cellular proportions and gene expression of specific cell types could better highlight cells and genes contributing to disease pathogenesis and identify therapeutic targets^8^. *In silico* deconvolution has been a promising approach to resolve the cellular composition at each measured spot. Most current ST deconvolution methods are reference-based, requiring a cell type-specific transcriptomic profile, usually generated from single-cell RNA-sequencing (scRNA-seq) experiments. With the reference profile, the cellular composition of each spot in the targeted ST dataset can be inferred. For example, RCTD^9^ learns the cell type profile from the reference dataset using a probabilistic model and predicts the cell type composition of a spot with maximum likelihood estimation. SpatialDWLS^10^ uses the scRNA-seq reference-derived signature to fit a dampened weighted least squares model to infer cell type composition. CARD^11^, as an autoregressive-based deconvolution method, combines cell type-specific expression information learned from the scRNA-seq reference with correlation in cell type composition across the tissue spots. Cell2location^12^ also borrows spatial information with a hierarchical Bayesian framework.

Despite the emerging number of scRNA-seq datasets, the desired reference profile may not be available for specific cell types or conditions^4^. The performance of these reference-based methods also highly depends on the quality of the reference profile, the sample processing techniques and the data processing procedures. The inferred cell types are also limited to those in the reference profile. Additionally, in some methods, the batch effects between the target ST dataset and the reference profile are not properly handled, resulting in inaccurate results^5^. Most importantly, by using a reference cell type-specific transcriptomic profile, it is assumed that the gene expression of each cell type is constant regardless of sample conditions such as disease status, ignoring the fact that there could be major differences in the cell type composition and cell type-specific gene expression across different sample conditions.

In contrast, reference-free methods do not require a scRNA-seq reference but rely on a list of marker genes for each cell type. STdeconvolve^13^, a Latent Dirichlet Allocation^14^ (LDA)-based reference-free method, decomposes the ST data into latent topics and simultaneously infers the topic-specific transcriptomic signature and topic compositions. Subsequently, by comparing the inferred topic-specific signature against known cell type marker genes with a gene set enrichment analysis (GSEA), each topic can be labelled with a cell type name. However, the estimated topics are usually redundant and cannot accurately capture cell populations with low abundance. Users may also obtain multiple topics corresponding to an abundant cell type and no topics for rare cell types. The estimates are also highly variable between runs, even when using the same dataset. CARD also offers a reference-free version (CARDfree^11^) that takes a list of marker genes as input to infer the cell type composition. However, the inferred cell types can be difficult to interpret and need to be further labelled with a GSEA.

To address these challenges, we present the deconvolution for <u>S</u>patial transcriptomics using <u>MAR</u>ker-gene-assisted <u>T</u>opic models (SMART), a reference-free deconvolution method based on semi-supervised topic models (**Figure 1**). In natural language processing, the topic models were used to identify the topic distribution from the words in a large number of unlabeled documents as well as the word frequencies within each topic. In the context of ST deconvolution, SMART simultaneously infers the cell type composition of the spots and the cell type-specific gene expression profile. Compared to unsupervised approaches such as STdeconvolve, which uses the marker information after the deconvolution process to label the latent topics, SMART directly incorporates marker gene information as prior knowledge during the topic inference procedures to guide cell type identification and, thus, improves the predictive accuracy and minimizes the variability. Using three datasets simulated from single-cell ST data and two real ST datasets, we demonstrate that SMART accurately estimates cell type composition and cell type-specific gene expression. Despite being a reference-free method, it outperforms some of the best-performing reference-based and reference-free methods for ST data^15–17^ when an ideal reference dataset is unavailable. Instead of a scRNA-seq reference, SMART uses a list of marker gene symbols for each cell type as the input. SMART also allows the inclusion of cell types with no marker gene information (“no-marker” cell type), which can be helpful in identifying novel cell types. The performance of SMART on cell subtypes can be augmented using a two-stage approach and the ability to obtain condition-specific estimation with a covariate model.

**Figure 1.**
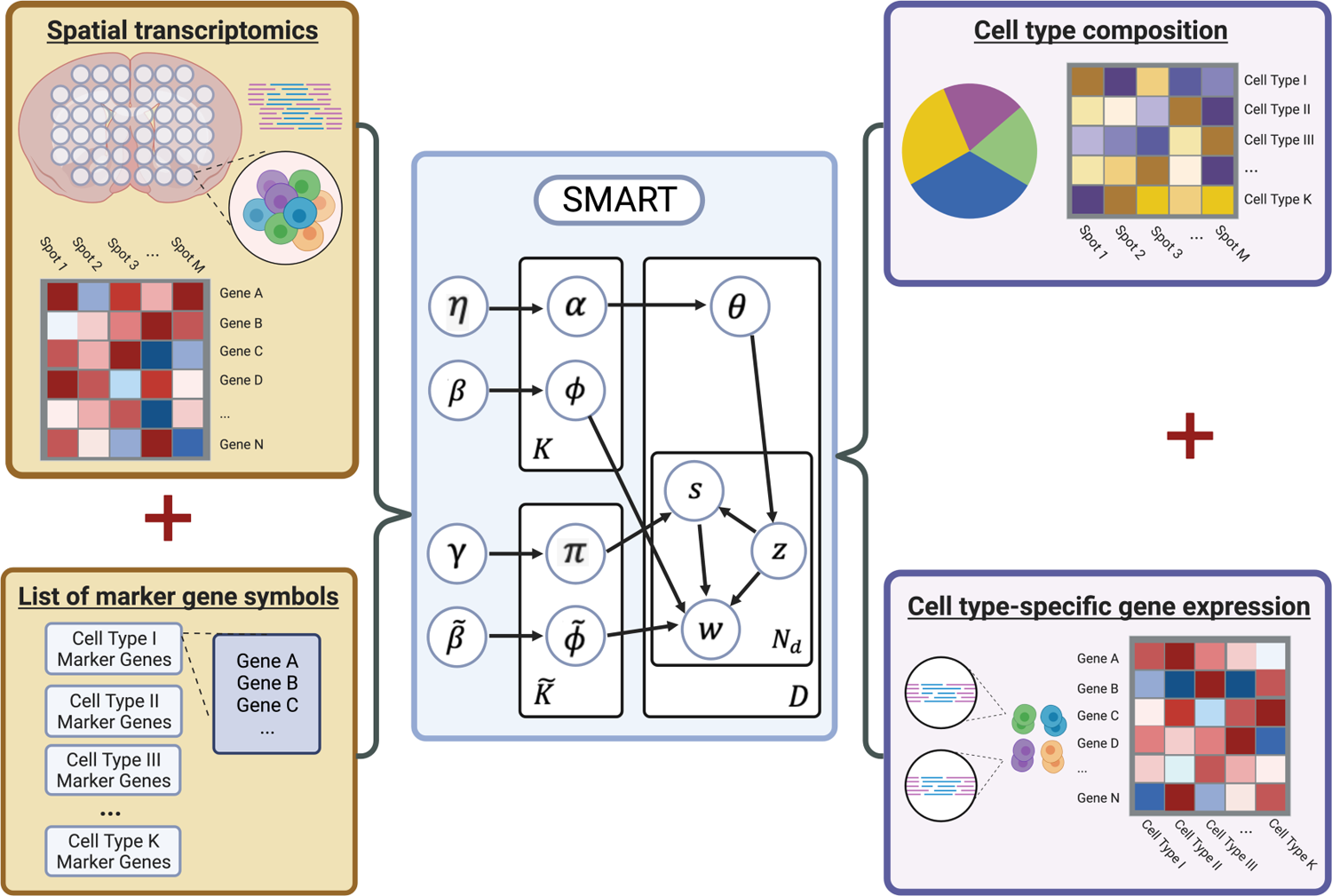
Overview of SMART. SMART takes the spatial transcriptomics data (a gene-by-spot matrix) and a list of marker gene symbols for each cell type as the inputs. Then, SMART uses a semi-supervised topic model to predict the cell type composition (a cell type-by-spot matrix) and the cell type-specific gene expression (a gene-by-cell type matrix) simultaneously. The model’s variables are described in Methods.

## Results

### SMART accurately predicted cell type composition in simulated ST data

To evaluate the performance of SMART, we used publicly available single-cell ST data in mouse kidneys (MK)^18^, which were profiled using the Vizgen Multiplexed Error-Robust Fluorescence in situ Hybridization^19^ (MERFISH) platform. The MK dataset contains the expression of 304 genes from 126,241 cells annotated to eight cell types. We simulated ST data by dividing the single-cell ST data of the MK dataset into 2474 spatially contiguous squares and aggregated the gene expression of cells within each square to mimic the spots of ST data (**Figure 2A**). The ground truth (GT) cell type proportion, cell type-specific gene expression and marker genes can be obtained accordingly from the simulated ST data and the original single-cell ST data. Then, we applied SMART to the simulated data along with the GT marker genes to simultaneously infer the cell type composition of each spot (**Figure 2B**) and the cell type-specific gene expression profile. We observed a strong correlation (>0.70) between the predicted cell type composition and the ground truth cell type composition in all cell types (**Figure 2C**).

**Figure 2.**
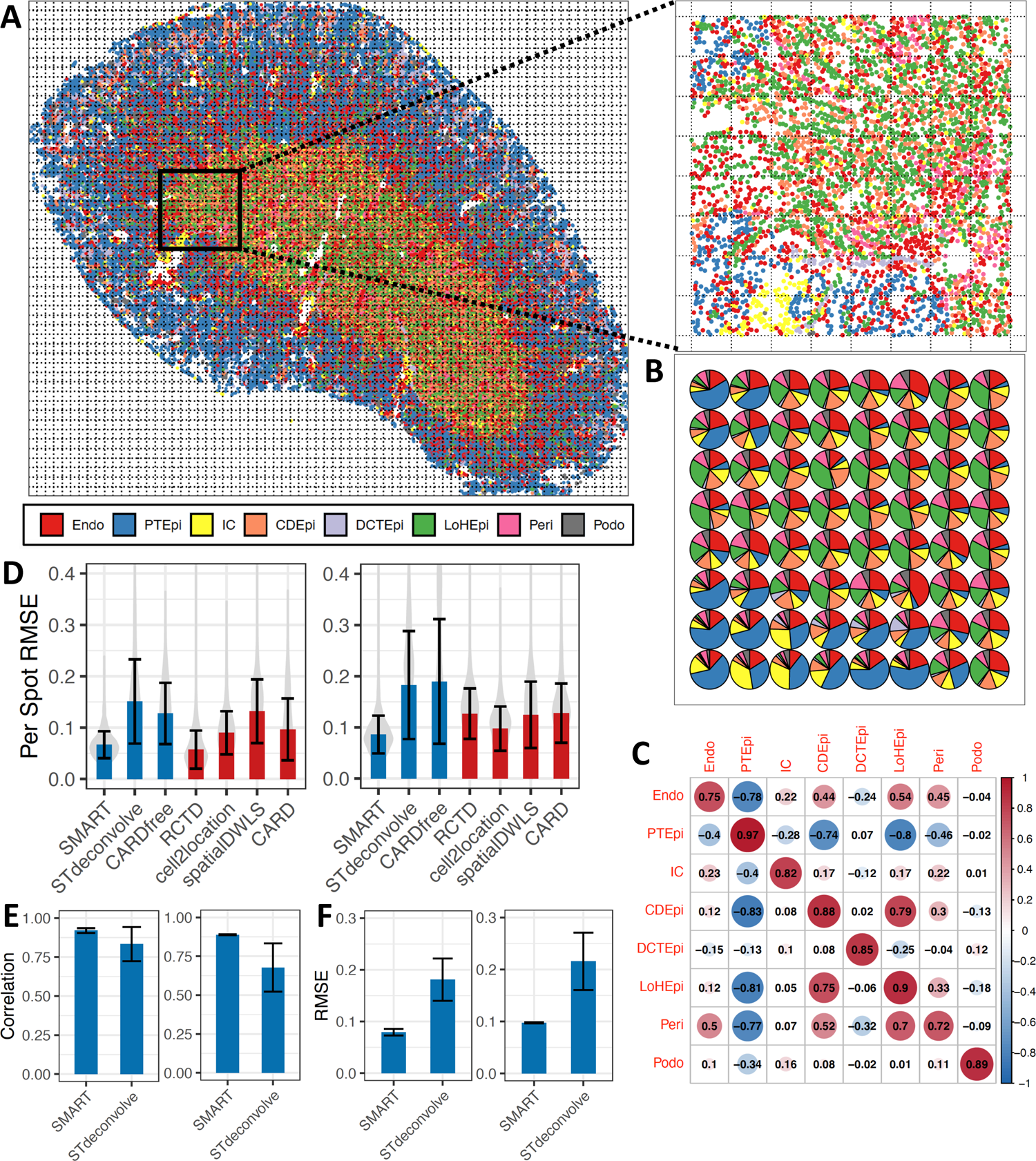
Performance evaluation of SMART using simulated ST data in mouse kidney. (A) The spatial image of the single-cell MERFISH data (left) and the zoom-in view of a selected area (right). (B) The SMART-predicted cell type composition of the selected area. (C) A heatmap showing the Pearson correlation coefficients between the predicted and the GT cell type proportions of each cell type. (D) The per-spot RMSE between the predicted and the GT cell type proportions using the GT (left) and the TMS markers/reference (right). Blue = reference-free methods; Red = reference-based methods. (E) The Pearson correlation between the predicted and the GT cell type proportions across all spots over 100 repeats using the GT (left) and the TMS markers/reference (right). (F) The RMSE between the predicted and the GT cell type proportions across all spots over 100 repeats using the GT (left) and the TMS markers/reference (right). Abbreviations: endothelial cell (Endo), epithelial cell of the proximal tubule (PTEpi), immune cell (IC), collecting duct epithelial cell(CDEpi), distal convoluted tubule epithelial cell (DCTEpi), loop of Henle epithelial cell (LoHEpi), pericyte (Peri), podocyte (Podo).

### SMART demonstrated superior performance to existing methods in realistic settings

To compare the performance of SMART against existing deconvolution methods, we next applied some of the best-performing reference-free (STdeconvolve, CARDfree) and reference-based methods (RCTD, CARD, cell2location, spatialDWLS) to the simulated MK dataset. For STdeconvolve and CARDfree, the GT marker genes were used to label the cell types; for reference-based methods, the original single-cell ST dataset used to simulate the ST data was used as the reference. We quantified the performance of each method with the Pearson correlation coefficient (PCC) and the root mean square error (RMSE) between the predicted and the GT cell type proportions of all spots across all cell types as well as the per-spot RMSE between the predicted and the GT cell type proportions of each spot. In the simulated MK dataset, SMART demonstrated better performance than most of the methods (PCC = 0.937, RMSE = 0.0715, mean per-spot RMSE = 0.0666, Diebold-Mariano P<0.001) except for RCTD^15–17^ (PCC = 0.955, RMSE = 0.0565, mean per-spot RMSE=0.0572) (**Figure 2D left**). However, using the original single-cell ST dataset as the reference profile to predict the ST dataset simulated from it will likely provide the best possible results for reference-based methods. Therefore, the reference-based methods were given a testing advantage over the reference-free methods. In reality, such an ideal reference profile with the same cell types and technical details rarely exists.

To examine the performance of the above methods in a more realistic situation where an ideal reference dataset is unavailable, we collected single-cell RNA-seq data in mouse kidneys from the Tabula Muris Senis (TMS) cell atlas^20^. The TMS single-cell dataset was used as the reference profile for reference-based methods to deconvolve the simulated MK dataset; the marker genes derived from the TMS dataset were used in reference-free methods. In this situation, SMART demonstrated the best performance over all the other methods (PCC = 0.921, RMSE = 0.0786, per-spot RMSE = 0.0860, Diebold-Mariano P<0.001, **Figure 2D right**). In both scenarios, SMART demonstrated the smallest variability in per-spot RMSE (**Figure 2D**).

### SMART demonstrated improved stability and interpretability than STdeconvolve

Similar to other reference-free methods based on generative models with sampling algorithms, the results of SMART and STdeconvolve may vary between runs according to the starting value. To assess the variability in performance, we performed 100 repeats of SMART and STdeconvolve on the MK dataset and examined the PCC and the RMSE between the predicted and GT cellular composition across all spots. We observed that SMART delivered more consistent results than STdeconvolve with less variability (**Figure 2E & F**). To further improve the stability, SMART provides the option to perform a user-specified number of repeats in parallel and average the results. Importantly, STdeconvolve identifies multiple latent topics for abundant cell types while identifying no topics for cell types that are less abundant. Out of the 100 repeats of STdeconvolve, 12 repeats identified only three cell types when using the GT marker genes in the GSEA to label the cell types; 58 repeats failed to identify more than five cell types; only four repeats were able to identify six cell types and no repeats identified all eight cell types. This indicates that the marker genes in SMART not only stabilize the deconvolution process but also ensure that we obtain an accurate estimate for each of the cell types.

### SMART demonstrated improved performance with a two-stage approach

Although marker genes can be shared across cell types in SMART, we recommend using marker genes with high specificity to achieve the best results. However, in some cases, marker genes can be very similar between cell types (i.e. monocyte, macrophage, and dendritic cells), especially those arising from the same lineage^21^, which makes it difficult to select specific marker genes. This ambiguity may lead to a drop in the accuracy of deconvolution results. To mitigate this limitation, we implemented a two-stage approach to improve the performance in predicting individual cell subtypes. In this two-stage approach, SMART was first applied to deconvolve the ST dataset into major cell types. Next, we extracted the gene counts explained by the cell type of interest. A second round of deconvolution was performed on the extracted gene counts to further decompose the cell type of interest to its subtypes. By separating the deconvolution process into two stages, any non-specific marker genes shared between the major cell types and those that were discarded in the first stage may be re-used in the second stage to discriminate the cell subtypes. In this manner, the selection of marker genes becomes easier for subtype identification compared to a one-stage approach that estimates all major cell types and cell subtypes during one run.

To illustrate, we collected a human non-small cell lung cancer (NSCLC) single-cell ST dataset^22^, which was profiled using the NanoString CosMx platform and simulated a ST dataset of 120 contiguous spots in the same manner (**Figure 3A**). The NSCLC single-cell dataset contains the expression of 960 genes from 32,272 cells. These cells were pre-annotated to ten major cell types or thirteen cell types with the inclusion of T cell subtypes (CD4+ T, CD8+ T, regulatory T) and dendritic cell (DC) subtypes (plasmacytoid DC and myeloid DC). In the two-stage approach, we first applied SMART to deconvolve the ST dataset into the major cell types (**Figure 3B**). Most cell types demonstrated a PCC > 0.90 between the GT and the predicted cell type composition (**Figure 3C**). Next, we extracted the gene counts explained by the T cells to further deconvolve them into T cell subtypes. The T cell subtype proportions obtained from the two-stage approach showed stronger correlations with the GT T cell subtype proportions compared to a regular one-stage approach (**Figure 3D top**). Users can also choose to include a cell type that is transcriptomically similar to the cell type of interest in the second round of deconvolution to recover potential incorrectly allocated gene counts. For example, to assist identification of DC subtypes, we included macrophages, which showed the greatest similarity to DCs in gene expression, during the second-stage deconvolution of DCs. By doing this, the correlation between the GT and predicted cell type proportion showed a dramatic increase for both pDCs and macrophages while staying similar for mDCs (**Figure 3D bottom**).

**Figure 3.**
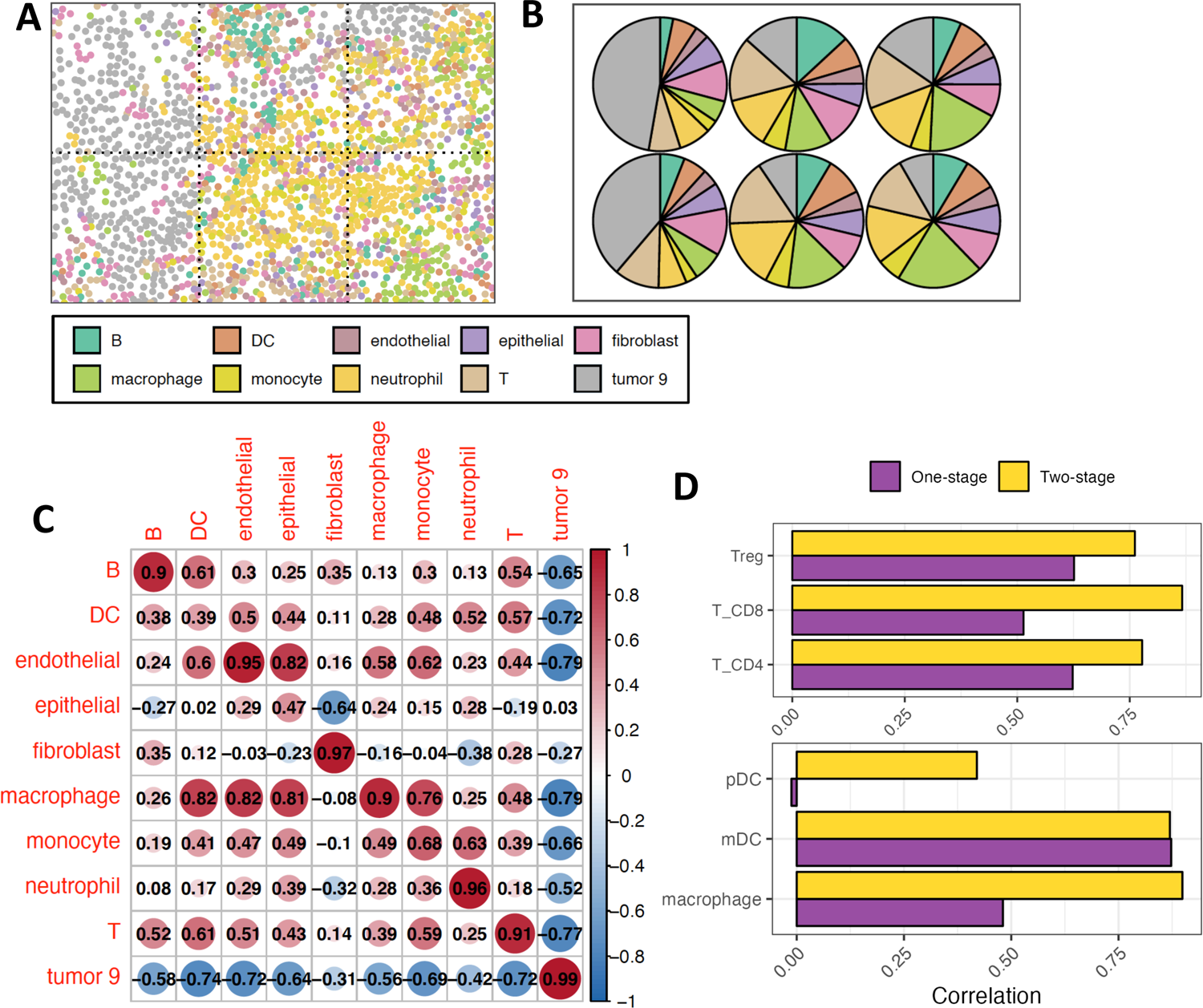
Demonstration of the two-stage approach of SMART. (A) An example field of view of the single-cell Nanostring CosMx data in human non-small cell lung cancer. (B) The SMART-predicted cell type composition of the example field of view. (C) A heatmap showing the Pearson correlation coefficients between the predicted and the GT cell type proportions of each cell type. (D) A comparison between the one-stage approach and the two-stage approach on predicting T cell subtypes (top) and dendritic cell subtypes (bottom).

### SMART identified condition-specific genes for each cell type with a covariate model

While reference-based methods assume that the cell type-gene expression is constant regardless of the sample conditions and that only the cell type composition can be changed, SMART respects the fact that cell type-specific gene expression can also vary between conditions. This was achieved by providing a covariate model allowing users to incorporate covariates to identify condition-specific genes for each deconvolved cell type. For demonstration, we simulated a ST dataset using a single-cell ST dataset on mouse hypothalamic preoptic area (MPOA) generated using the MERFISH platform^23^ (**Figure 4A**). The MPOA dataset contains the expression of 135 genes from 49,138 cells of a female mouse. We first validated the performance of the base model on the simulated MPOA dataset (**Figure 4B**) and observed a correlation of > 0.75 between the GT and the predicted cell type proportions in all of the cell types except microglia (r=0.65, **Figure 4C**). Next, to evaluate the performance of the covariate model, we re-simulated the ST dataset by including the data on another male mouse from the MPOA single-cell dataset. Then, we applied the covariate model using sex as the covariate and obtained the cell type-specific gene expression for the female mouse and the male mouse. Consistent with the literature, we observed that in excitatory neurons, Brs3 was up-regulated in the female mouse (**Figure 4D top**); in inhibitory neurons, Esr1 was up-regulated in the female mouse while Sytl4, Cyp19a1 and Greb1 were up-regulated in the male mouse^23^ (**Figure 4D bottom**). SMART then incorporates the differences in their cell type-specific gene expression profile with GSEA, identifying pathways that were enriched due to sex differences. For example, we found that ion, inorganic molecular entity, and salt transmembrane transporter activity were up-regulated in inhibitory neurons of the female mouse in comparison with those of the male mouse (Benjamini-Hochberg false discovery rate < 0.05).

**Figure 4.**
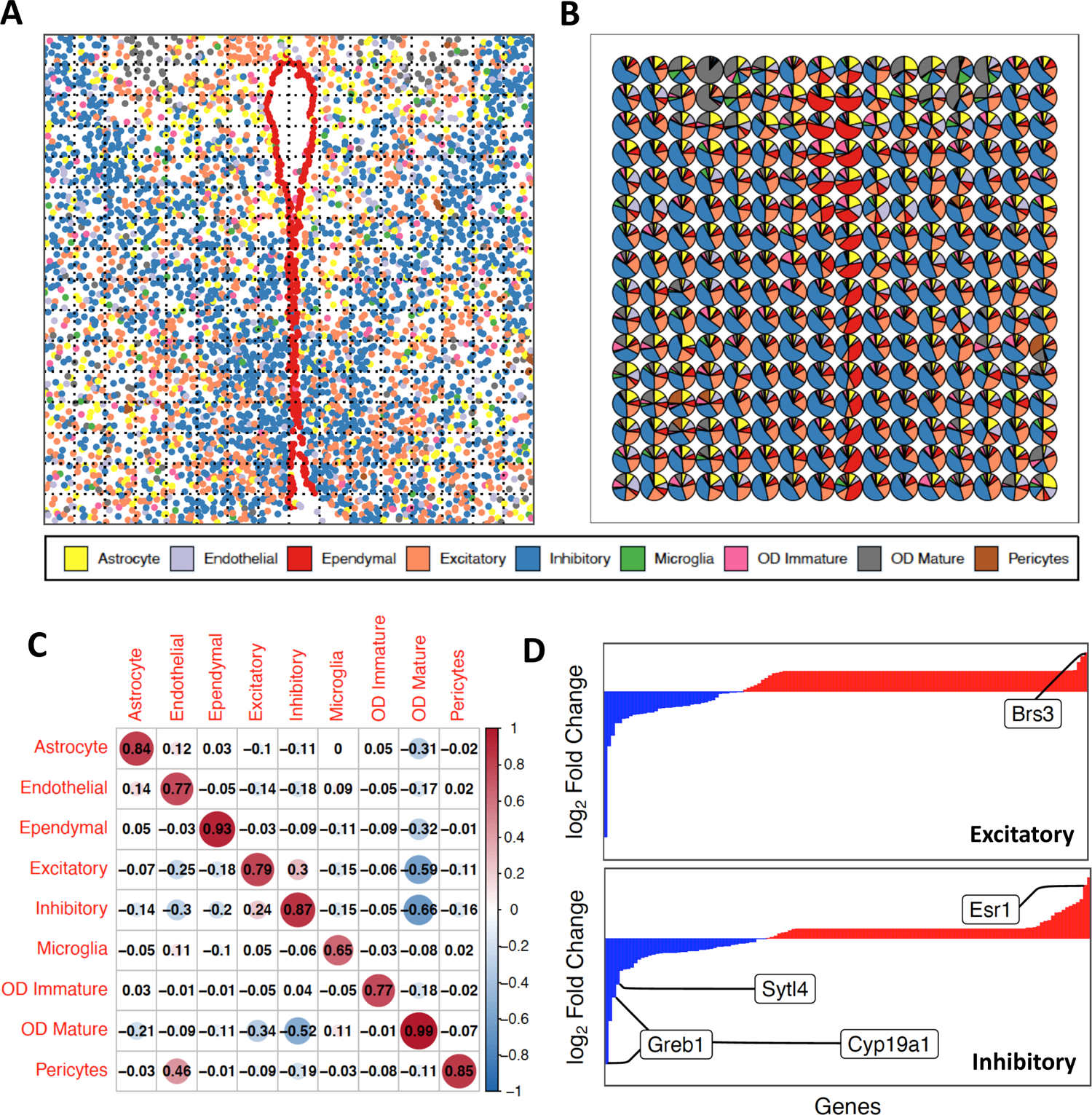
Demonstration of the covariate model of SMART using simulated data in mouse hypothalamic preoptic region. (A) Spatial image of the single-cell MERFISH data. (B) The SMART-predicted cell type composition. (C) A heatmap showing the Pearson correlation coefficients between the predicted and the GT cell type proportions of each cell type. (E) The log2 fold change of gene expression between the female mouse and the male mouse in the excitatory neurons (top) and the inhibitory neurons (bottom). Red bars = genes up-regulated in the female mouse; Blue bars = genes up-regulated in the male mouse.

### SMART is compatible with diverse ST platforms

To validate the performance of SMART on real ST datasets, we applied SMART to a mouse brain ST dataset^24^, which was profiled using the 10X Visium platform (**Figure 5A**). Depending on the tissue type, the Visium platform usually contains 1-10 cells per spot^3^. With marker genes identified from the Mouse Brain Atlas^25^, we deconvolved the mouse brain ST dataset into seven known cell types as well as an unknown cell type (**Figure 5B**). SMART successfully identified cell types in brain regions such as fiber tracts, ventricles, cortex, thalamus and hypothalamus. As expected, the oligodendrocytes were predicted to be highly enriched in fiber tracts compared to non-fiber tract regions (t-test P<0.05, **Figure 5C & D**); neurons were highly enriched in non-fiber tract regions as opposed to fiber tracts (t-test P<0.05, **Figure 5E**). Similarly, we observed a high predicted proportion of ependymal cells in ventricular regions compared to non-ventricular regions (t-test P<0.05, **Figure 5F & G**). Interestingly, the unknown cell type we obtained may correspond to a cell type in the medial habenula region of the mouse brain (**Figure H**). This suggests that SMART, as a reference-free method, may help identify novel cell types without marker genes.

**Figure 5.**
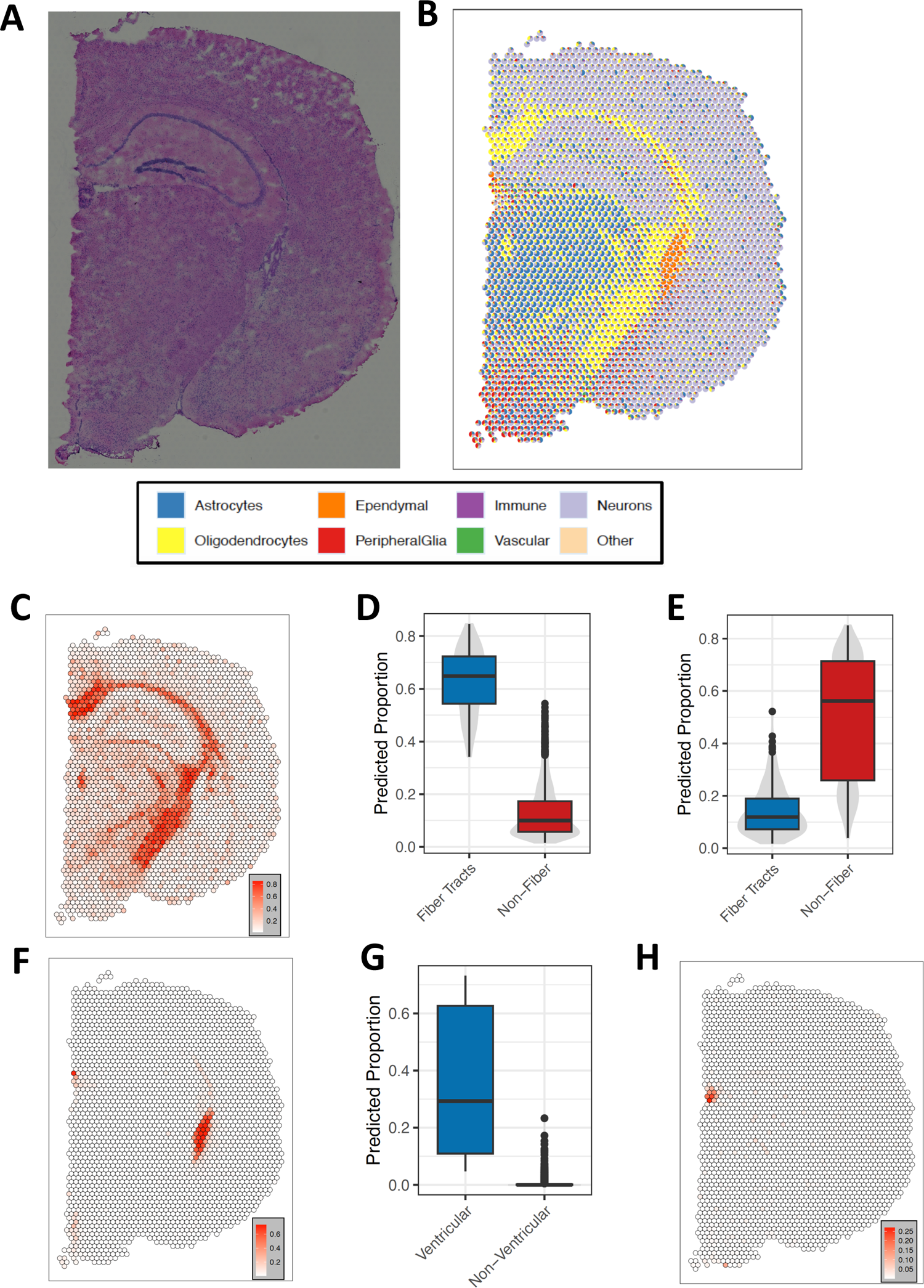
Application of SMART on a mouse brain ST dataset profiled using the 10X Visium platform. (A) Histology staining image of the tissue. (B) SMART-predicted cell type composition. (C) The predicted proportion of oligodendrocytes in each spot. (D) A comparison of the predicted proportion of oligodendrocytes in the spots of the fiber tract region and the spots of the non-fiber tract region. (E) A comparison of the predicted proportion of neurons in the spots of the fiber tract region and the spots of the non-fiber tract region. (F) The predicted proportion of ependymal cells in each spot. (G) A comparison of the predicted proportion of ependymal cells in the spots of the ventricular region and the spots of the non-ventricular region. (H) The predicted proportion of the unknown cell type in each spot.

In addition to the mouse brain ST dataset, we also validated the performance of SMART on an ST dataset generated from human pancreatic ductal adenocarcinoma sample^26^ using microarray slides (**Figure 6A**). With marker genes derived from its sample-matched scRNA-seq dataset generated using the inDrop platform, SMART identified sixteen cell types across four distinct tissue regions labelled based on histology staining (**Figure 6B-D**). As expected, we observed a higher predicted proportion of ductal cells in the ductal region than in the non-ductal regions (t-test P<0.05, **Figure 6E**). Also, we observed a higher predicted proportion of acinar cells in the pancreatic region (t-test P<0.05, **Figure 6F**) and cancer clone cells in the cancerous region (t-test P<0.05, **Figure 6G**).

**Figure 6.**
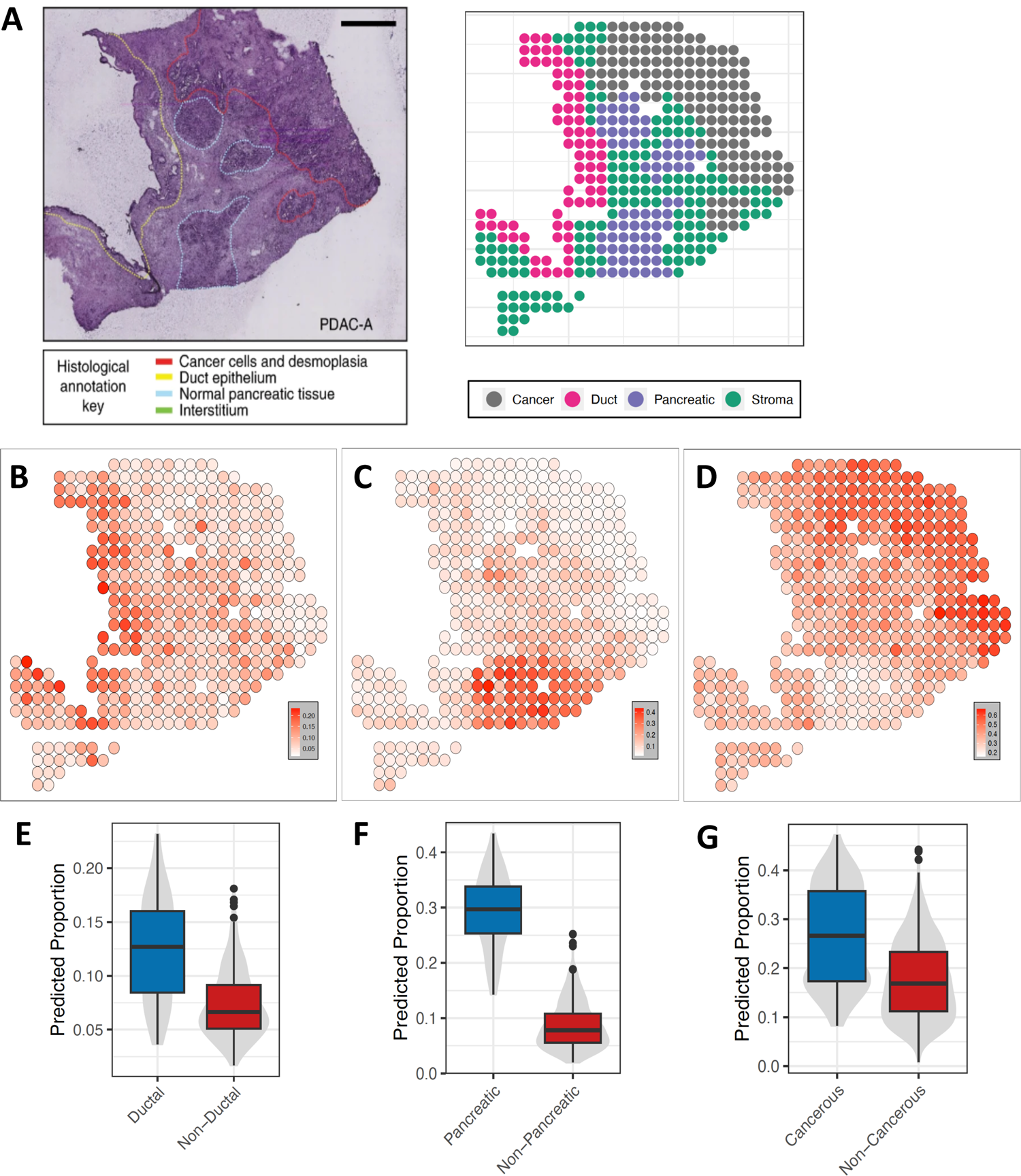
Application of SMART on a pancreatic ductal adenocarcinoma ST dataset. (A) Histology staining image of the tissue (left) and the regions annotated by histologists from the original study (right). (B) The predicted proportions of ductal cells. (C) The predicted proportions of acinar cells. (D) The predicted proportions of cancer clone cells. (E) A comparison of the predicted proportion of ductal cells in the spots of ductal region and the spots of the non-ductal region. (F) A comparison of the predicted proportion of acinar cells in the spots of the pancreatic region and the spots of the non-pancreatic region. (G) A comparison of the predicted proportion of cancer clone cells in the spots of the cancerous region and the spots of the non-cancerous region.

### Factors affecting the performance of SMART

Finally, to assess factors that may affect the results of SMART, we first examined if the total number of spots affects the deconvolution performance. With the simulated MK dataset, we randomly selected 50, 100, 300, 500 and 1000 spots and applied SMART. The results show that interrogation of more spots did not lead to a significant decrease in the per-spot RMSE (t-test P>0.05, **Supp. Figure 1A**).

Next, we examined if the number of cells in each spot has any effect on SMART. With technological improvement, the resolution has become higher in recent ST platforms^27^. Using the NSCLC dataset, we re-simulated the ST data by decreasing the size of the contiguous squares to contain, on average, 269, 68 and 17 cells per spot to mimic the scenario that higher-resolution platforms usually have more spots with a smaller tissue coverage area in each spot. Interestingly, while the change in per-spot RMSE is subtle, it decreased with a higher number of cells per spot (t-test P<0.001, **Supp. Figure 1B**), suggesting that SMART may perform better on ST platforms with a lower resolution.

Most importantly, since the selection of input marker genes can affect the performance of SMART, we examined how the number of marker genes affects the results of SMART by using at most 3, 5, 10 and 15 marker genes per cell type. As anticipated, SMART demonstrated a lower per-spot RMSE as the number of marker genes increased (t-test P<0.001, **Supp. Figure 1C**). This suggests that including more marker genes improves the performance of SMART, assuming the quality of the marker genes. The largest decrease in mean per-spot RMSE occurred between having five marker genes and having ten marker genes (7.13% decrease). These data suggest that having approximately ten marker genes per cell type may efficiently improve the performance of SMART.

## Discussion

Using multiple simulated and real ST datasets, here, we demonstrated that SMART accurately captures the cell type composition and cell type-specific gene expression of ST data across various ST platforms, even in comparison to some of the best-performing reference-based methods^15–17^. Using the MK simulated dataset, we showed that in an ideal situation, where we used the original single-cell dataset as the reference to deconvolve the ST dataset simulated from it, SMART showed comparable performance to the reference-based methods. Although methods such as RCTD may show a slightly better performance in an optimal setting, such conditions rarely exist in the real world. In a more realistic circumstance where we used an external reference dataset, SMART shows significantly lower error compared to reference-based methods, indicating that SMART, as a reference-free tool, provides more accurate results in real-world settings. Without the need to properly process a reference dataset, SMART minimizes the impact of batch effects and is also easier to use compared to most reference-based methods. The results from the NSCLC dataset showed that the two-stage approach might help to further improve SMART’s performance in decomposing cell subtypes by optimizing the use of marker genes and by recovering falsely allocated gene counts. Finally, we used the MPOA dataset to demonstrate the ability of SMART-covariate to estimate the condition-specific gene expression profile for each cell type by including the condition as a covariate. An important assumption in the reference-based methods is that the cell type-specific gene expression is constant across sample conditions, and only cell type compositions differ. However, this assumption is frequently violated in the real world as gene expression in samples is modified by disease and treatment, leading to inaccurate results. SMART surmounts this limitation by enabling the inclusion of covariates that can capture the impact of disease status and phenotypes. A recent method Celloscope also uses a marker-gene-driven probabilistic model to perform deconvolution^28^; however, it does not provide a two-stage approach nor a covariate model. In addition, the ability to include cell types without marker genes assists in identifying novel cell types, as shown in the analysis of the 10X Visium mouse brain dataset. With the two real ST datasets, we showed the compatibility of SMART on high-resolution platforms such as the 10X Visium as well as lower-resolution platforms such as the one used in the PDAC dataset.

SMART allows shared marker genes between cell types; however, using markers specific to cell types usually provides better results. While obtaining high-quality marker genes can be challenging, SMART only requires a small number of marker genes. As illustrated with the NSCLC dataset, we can obtain a good to excellent estimation with as few as three marker genes per cell type. The results can be improved efficiently by using approximately ten marker genes per cell type.

There are also limitations to SMART. Firstly, all the spots are assumed to be independent in SMART instead of borrowing spatial information from the adjacent spots. However, SMART does include a gene weighting scheme to borrow gene abundance information from all spots to prevent certain genes from dominating a cell type. Moreover, the selection of input marker genes is critical to optimize the performance of SMART. To make the tool more accessible, SMART provides several pre-defined lists of marker genes for common tissue types.

In summary, we present SMART as a reference-free deconvolution method for spatial transcriptomics without needing a scRNA-seq reference profile. By incorporating marker genes for the cell types as guidance, SMART showed improved accuracy, stability and interpretability even when compared with some of the best-performing reference-based methods. With the two-stage approach and the covariate model, SMART shows an advantage in discriminating the cell subtypes and studying biological perturbations. Ultimately, we believe that SMART will be a powerful tool to unravel the tissue heterogeneity and identify potential therapeutic targets at a single-cell-type resolution with spatial information.

## Methods

### Overview of SMART

SMART builds on the keyword-assisted topic models (keyATM)^29^, which are semi-supervised topic models that integrate prior knowledge to guide the formation of topics. By including a small number of keywords for each topic prior to model fitting, keyATM accurately infers the proportion of topics within each document and the word frequencies within each topic.

In the context of cell type deconvolution in SMART, the spots correspond to the documents, the genes correspond to the words, and the cell types correspond to the inferred topics. A small number of marker genes for each cell type (keywords) were used as prior knowledge to help infer the cell type proportions (topic proportions within each document) and the cell type-specific gene expression (word frequencies within each topic) in the form of relative gene frequencies.

The ST data is represented as a *N* × *D* matrix with *N* genes and *D* spots. The total number of RNA molecules in each spot *d* is *N*_d_. We use *W*_di_ to represent the *i*th gene in spot *d*. KeyATM is a generative model based on a mixture of two Dirichlet distributions, one for the marker genes and one for other non-marker genes. A key assumption is that the marker genes should have a higher frequency in a given cell type than the non-marker genes. The data generation process is as follows:

1. Suppose there are a total of *K* cell types and the first K^∼^ of them are cell types provided with marker genes.
2. For each gene *i* in spot *d*, we draw the cell type variable *Z_di_* from the topic distribution:

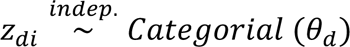

θ*d* represents the cell type proportions within each spot *d*.
3. If the sampled cell type *k* is a “no-marker” cell type, we draw the gene *w_di_* from the gene distribution:

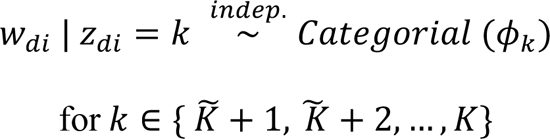

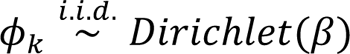 represents the gene frequencies within cell type *k*.
4. If the sampled cell type contains gene markers, we first draw a Bernoulli random variable *s_di_* with success probability G_’_ for gene & in spot *d*

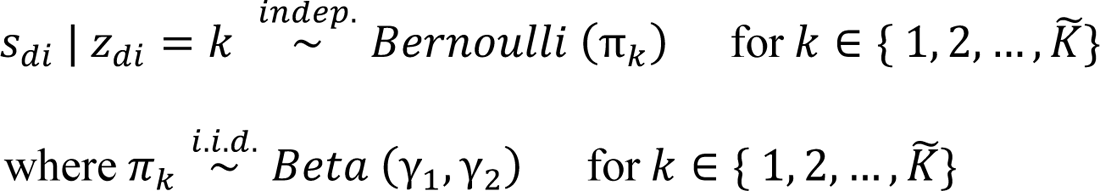

If the variable equals 1, gene *w_di_* is drawn from the gene frequencies for cell type *k* with marker genes 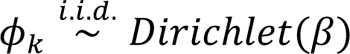; if the variable equals 0, *w_di_* is drawn from the gene frequencies for cell types without marker genes Φ_k_. That is,

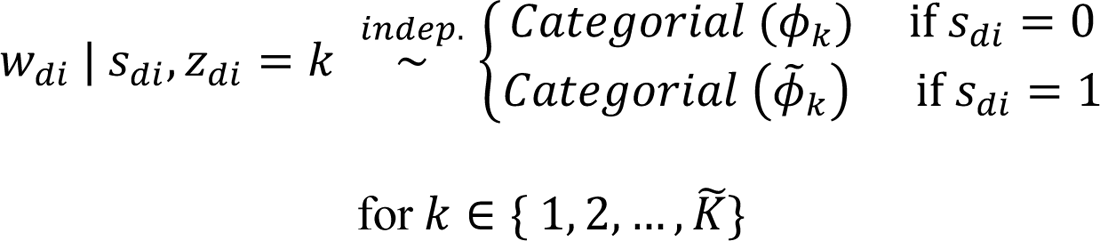

β and β^∼^ are hyperparameters to make the prior means for the frequency of marker genes higher than those of non-marker genes.
5. And,

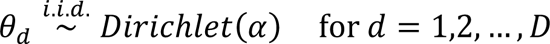

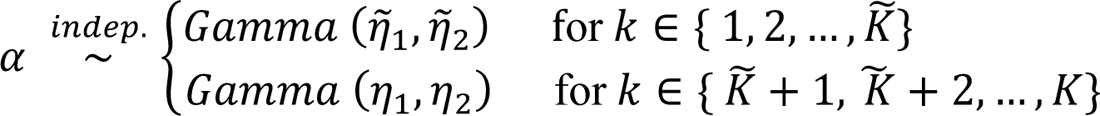

η^∼^_1_ and η^∼^_2_ were set to sample from larger values so that the spots are more dominant by cell types with marker genes.

**Figure 1** shows a graphic representation of this generative process. By integrating out the latent variables (θ, φ, φ^∼^, π), keyATM uses a collapsed Gibbs sampling algorithm to sample from their posterior distribution. It also uses an inverse gene frequency weighting strategy to help prevent highly expressed genes from dominating the inferred cell types. The resulting 5 matrix represents the cell type proportions for each spot; Φ and Φ^∼^, combining, represents the cell-type specific gene expression.

### The two-stage approach

To better estimate the composition of cell subtypes of a major cell type *k*, we take SMART one step further to perform a second round of deconvolution as follows:

1. Perform the first round of deconvolution and during deconvolution.
2. Calculate the total library size (sum of the gene counts) of each spot *d*_T_
3. Calculate the library size for each cell type of each spot *d*_K_ with *d*_T_θ, where θ is the cell type composition matrix.
4. Calculate the gene counts for cell type *k* of with *d*_k_φ_k_, where *d*_k_ is the library size for cell type *k* of each spot and φ_k_ is the relative gene frequency for cell type *k*.
5. Identify another cell type *m* with the highest similarity by calculating the PCC or the Euclidian distance between the relative gene frequency of the cell types. This cell type *m* can also be user-defined.
6. Calculate the gene counts for cell type *m* as in step 4).
7. Perform the second round of deconvolution on the sum of the gene counts for cell type *k* and *m*.

SMART includes a “no-marker” cell type in both stages to represent any data that cannot be explained by the cell types with marker genes.

### The SMART-covariate model

SMART-covariate extends the base model and builds on the keyATM covariate model. Instead of step 5) in the base model, the covariate model uses the following cell type distribution:

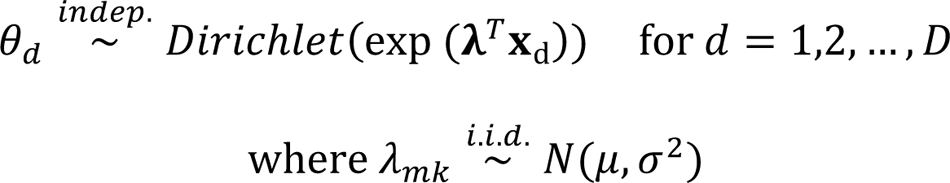

where **x**_d_ is an *M*-dimensional covariate matrix for each spot *d*. λ is an *M* × *K* matrix of coefficients and λ_mk_ is the (*m, k*) element of λ.

### Simulating ST data

To mimic the spots of ST data, we collected pre-annotated single-cell resolution ST data and cut the spatial image into contiguous squares. To start with, the bottom and left edge of the bottom-left square was aligned with the bottom and the left edge of the spatial image. Then, the squares were created by drawing boundaries in increments of a selected value based on the datasets until they reached the top and right edge of the spatial image. Any squares with less than two cells were removed from the simulated dataset. The squares overlapping with the spatial image’s edges were also removed. The gene counts of cells within the coordinate of each square were aggregated together to obtain the gene expression of this simulated spot. The GT cell type proportions of each spot can be calculated by counting the number of cells of each cell type in each square. The GT cell type-specific gene expression can be obtained by averaging the gene expression of cells of the same cell type in the original single-cell ST dataset. Finally, the GT gene markers for each cell type can be obtained through a differential expression analysis between one cell type and the rest. The differential expression analysis was performed with a Wilcoxon rank-sum test, and the Benjamini-Hochberg procedure was used to correct for multiple hypotheses testing. A false discovery rate < 0.05 was used as the threshold for significant gene markers. The resulting markers were also filtered by fold changes to select markers of high confidence. The gene markers were further pruned to keep marker genes specific to each cell type.

### Deconvolution using SMART and other existing methods

In SMART, the input ST data could be either gene counts or un-transformed normalized gene expression rounded to integers. A list of gene symbols for marker genes of each cell type was used as the supplemental input to guide cell type inference. By including the marker genes prior to deconvolution, the resulting cell type proportions and cell type-specific gene expression were automatically labelled with cell type names, improving the results’ interpretability. In addition to cell types with marker genes, SMART also allows the inclusion of unknown cell types without specifying any marker genes, which may be helpful in identifying novel cell types. The GSEA in SMART-covariate was implemented using the R package “fGSEA”^30^.

STdeconvolve, as an unsupervised reference-free method, used the ST data as the only input. The resulting cell type proportions and cell type-specific gene expression matrices contained no cell type names. These unlabeled cell types were subsequently annotated with a name through a GSEA by comparing the inferred cell type-specific gene expression profile against the marker genes using the R package “liger”^31^.

CARDfree requires marker genes as the input in addition to the ST data. In all the analyses, the same gene markers used in SMART were used as the input for CARDfree. The inferred cell types, however, were not annotated with any cell type names. The same gene set enrichment analysis approach used in STdeconvolve was applied to label the cell types with cell type names.

The reference-based methods (RCTD, cell2location, spatialDWLS, CARD) were performed using the recommended settings according to the guidelines on their websites. These methods require a reference single-cell RNA-sequencing dataset as the input instead of a list of gene symbols used in reference-free methods.

## Data availability

Data from the MK dataset is available for download from https://figshare.com/projects/MERFISH_mouse_comparison_study/134213. Data from the MPOA is available at: doi: https://datadryad.org/stash/dataset/10.5061/dryad.8t8s248/. The 10X Visium coronal section of the mouse brain dataset is available for download at https://www.10xgenomics.com/resources/datasets/mouse-brain-section-coronal-1-standard-1-1-0

The PDAC data were obtained from sample A of the PDAC dataset and are available at the Gene Expression Omnibus (accession number GSE111672).

## Code availability

SMART is available as an R package on GitHub at https://github.com/yyolanda/SMART.

## Contribution of authors

All authors contributed to project design, data analysis and results interpretation. CY contributed to the implementation of the software package. All authors contributed to the writing of the manuscript.

---

**Supp. Figure 1.**
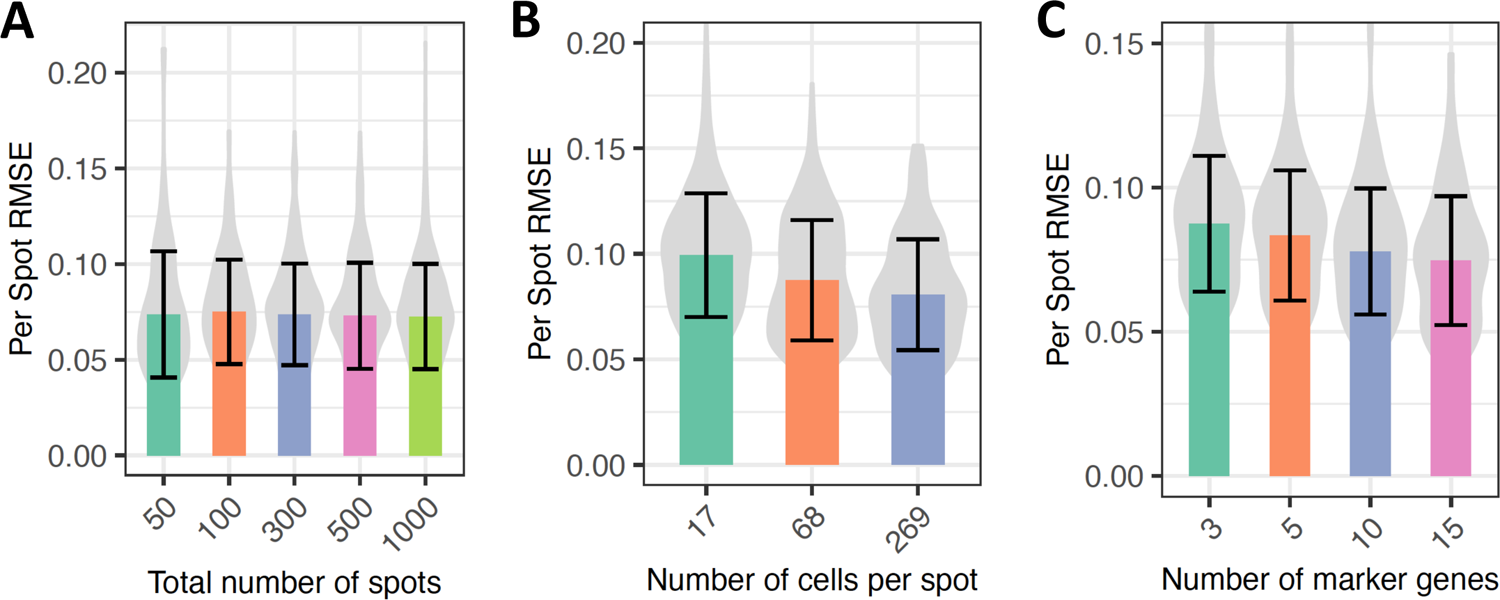
Factors that may affect the performance of SMART. (A) The number of total spots. (B) The number of cells on average within each spot. (C) The number of marker genes.

